# Nuclear rerouting of paracrine Fgf3 in source cells represses target genes to pattern morphogen responses

**DOI:** 10.1101/2025.07.24.666518

**Authors:** Sevi Durdu, Murat Iskar, Alicia Roig-Merino, Elisa Gallo, Ezgi Karaca, Andreas Kunze, Erika Donà, Dirk Schübeler, Peer Bork, Darren Gilmour

## Abstract

Morphogen gradients direct tissue patterning by inducing dose-dependent transcriptional responses, yet ligand-producing cells often respond differently from their neighbors. Using the zebrafish lateral line organogenesis model, we uncover a cell-autonomous role for the paracrine ligand Fgf3. Transcriptomic profiling and quantitative single-molecule imaging identify target genes, including the chemokine scavenger *cxcr7b*, whose expression decreases both when FGF receptor signaling is inhibited and when Fgf3 is overexpressed. High-resolution live imaging reveals nuclear accumulation of Fgf3 in producing cells, whereas neighbors receive only extracellular ligand, a feature also observed in other embryonic tissues. Mosaic gain-of-function and nanobody-mediated degradation demonstrate that the nuclear pool of Fgf3 autonomously represses specific targets without impairing canonical receptor signaling. Structure-guided comparative assays indicate nuclear targeting as a latent property of several paracrine FGFs. Dual secreted–nuclear functionality of FGF ligands may represent an intrinsic symmetry-breaking mechanism during organogenesis.

## Introduction

Multicellular development depends on the regulated expression of genes that drive the differentiation of cell groups into tissues and organs of defined form and function. This spatiotemporal regulation is provided by potent secreted signals termed morphogens ^1–3^. Fibroblast growth factors (FGFs) are well-studied morphogens that control diverse developmental processes, including early tissue specification, somitogenesis, and proximodistal limb patterning ^4–7^. Studies in these and other model systems have provided key mechanistic insights into how morphogens generate complex spatiotemporal patterns of gene expression ^8–11^. At the molecular level, secreted FGF ligands typically activate tyrosine kinase receptors, which in turn phosphorylate Ets transcription factors in the nucleus to enhance positive pathway targets and repress negative ones ^7,12–15^. At the tissue scale, expression patterns are thought to arise primarily from the graded distribution of FGF ligands and the differential dose responses of target genes, each regulated at defined morphogen thresholds ^11,16^. These mechanisms, however, can be further modulated by cell-specific features that remain less well understood. For example, cells at the source of morphogen gradients are exposed to the highest ligand levels, yet in many contexts respond less strongly than their immediate neighbors ^3,17,18^. In some cases, this difference reflects reduced receptor expression in ligand-producing cells, leading to attenuated signaling ^19^. More generally, however, the mechanisms that distinguish morphogen source cells from responders in differentiating tissues remain elusive.

To investigate the roles of FGF signaling in gene regulation, we utilized the development of posterior zebrafish lateral line primordium (pLLP), a tractable in vivo model for studying dynamic tissue patterning at cellular resolution^20–22^. In this system, localized sources of FGF ligand expression direct the orderly assembly and deposition of rosette-like mechanosensory organs, termed neuromasts, by a migrating collective of epithelial cells^23–25^. The range of FGF action within the pLLP is restricted by the expression of extracellular matrix-associated molecules, such as HSPGs and Anosmin1, and the formation of multicellular lumina that concentrate the secreted ligand at the center of each neuromast progenitor ^26,28,29^. While these mechanisms explain how FGF acts locally to promote neuromast assembly, it remains unclear whether and how FGF morphogens drive cell type diversification within these differentiating organs.

Here, we investigate the impact of FGF signaling on gene expression patterning in the pLLP using RNA profiling combined with in vivo subcellular analyses. We show that tissue-specific upregulation of Fgf3 leads to transcriptional repression of a subset of genes that otherwise behave as bona fide positive targets of FGF receptor signaling, including the chemokine receptor *cxcr7b*. In line with this unexpected finding, mosaic expression of Fgf3-GFP demonstrates that repression of *cxcr7b* is autonomous to source cells and can be mediated by non-secreted forms of the ligand. Targeted depletion experiments further confirm that this activity depends on a nuclear-localized pool of Fgf3. Beyond contributing to the growing literature on proteins with proposed dual functionality^27^, these data support a general mechanism that distinguishes morphogen source cells from responders.

## Results

To obtain a comprehensive view of the gene regulatory targets of FGF signaling in the posterior lateral line primordium (pLLP), we manipulated FGF activity using two approaches. First, we applied the small-molecule antagonist SU5402 to inhibit the kinase domain of FGF receptor 1 (Fgfr1), at a concentration previously shown to block FGF-dependent organ differentiation and target gene expression in this tissue (“Fgfr1-inhibited”). Second, we employed an inducible FGF ligand expression system that we had previously validated as a tunable enhancer of FGF signaling (“Fgf3-induced”)^28,29^. Treated pLLP cells were then isolated with high specificity by fluorescence-activated cell sorting (FACS), using expression of two fluorescent transgenic reporters, cxcr4b:nlsTomato^30^ and cldnb:lyn-GFP^25^, as selection markers (Figure S1). We next established the gene expression profiles of wild-type, Fgfr1-inhibited, and Fgf3-induced pLLPs in six replicates. Profiling and bioinformatic analysis revealed a comprehensive catalog of pLLP-specific genes with distinct responses to FGF signaling (Figure 1A, Table S1).

**Figure 1.**
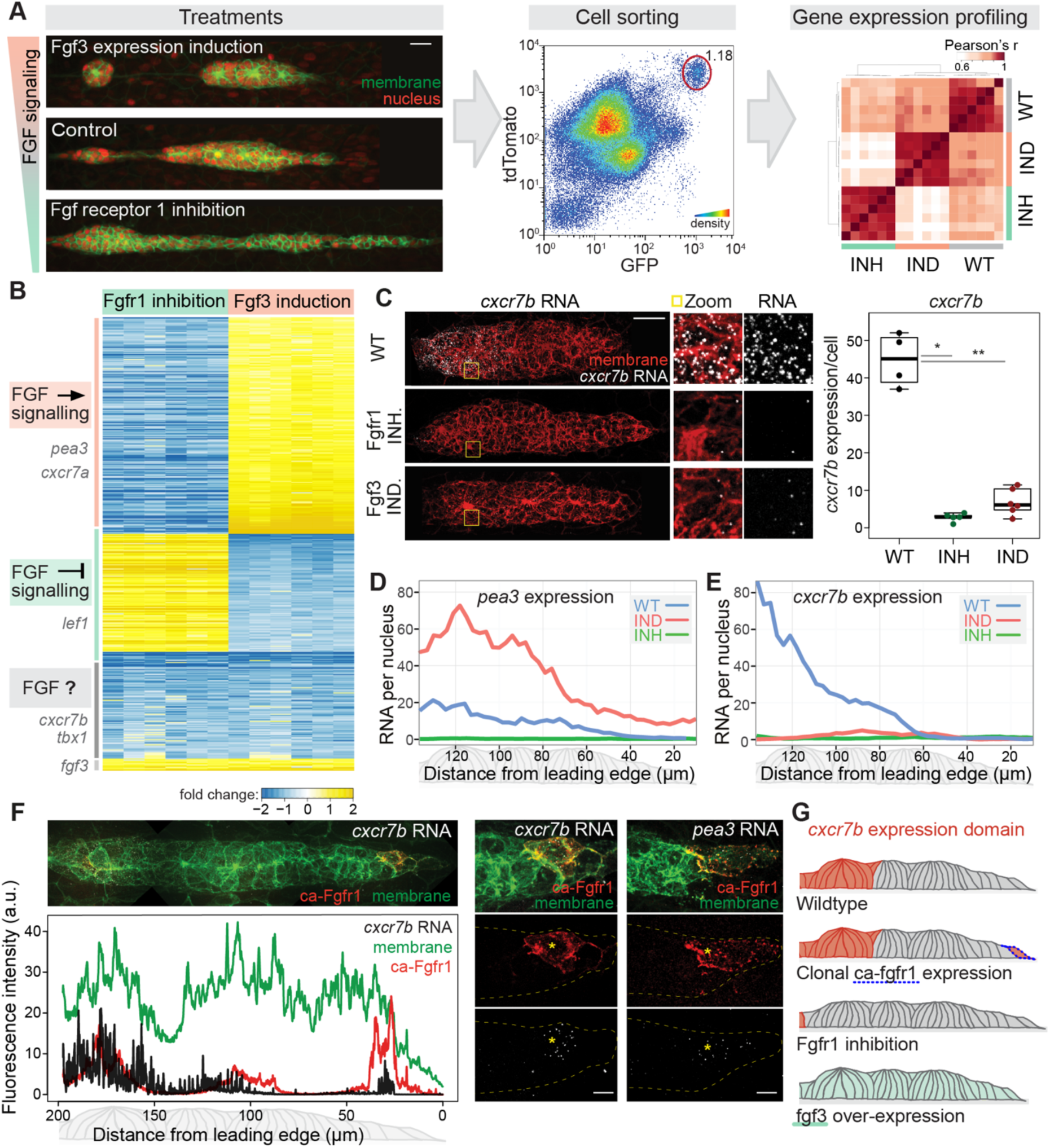
Gene expression profiling following FGF signaling perturbations uncovers atypical regulation of a subset of target genes in the zebrafish lateral line. **A.** Gene expression profiling pipeline. Left: Images of lateral line primordium upon manipulation of FGF signaling levels through FGF ligand induction (*fgf3* overexpression) and FGF receptor inhibition (Fgfr1 kinase domain inhibition with SU5402) on cldnb:lyn-GFP (green membrane), cxcr4b:nlsTomato (red nucleus) transgenic lines alongside controls. Middle: Isolation of lateral line cells from the rest of the embryo by FACS using fluorescent markers and scatter features. Red circle indicates the pLLP population (see Figure S1). Right: Gene expression profiling of 6 biological replicates with 3 treatment conditions; the graph shows hierarchical clustering of the 18 samples. **B.** Heatmap of differentially expressed genes in response to both Fgfr1 inhibition and Fgf3 induction (blue: downregulation, yellow: upregulation). Left: Anti-correlated genes are dose-responsive genes oppositely affected by Fgf3 induction and Fgfr1 inhibition (e.g., *pea3* and *lef1*), whereas correlated (dose-independent) genes respond in the same direction (e.g., *cxcr7b*). Gene expression profiles recapitulate previously documented responses to FGF signaling inhibition, including *pea3* as a positive and *lef1* as a negative target of FGF signaling. C. smFISH-based quantification of *cxcr7b* expression in pLLP across wild-type (WT), Fgf3 induction (IND), and Fgfr1 inhibition (INH) conditions. Zoomed regions are boxed. Boxplot shows *cxcr7b* transcript counts per cell, per primordium (n = 4, 5, 6). **D–E.** smFISH-based expression profiles of the canonical FGF signaling target *pea3* and the dose-independent target *cxcr7b* along the pLLP in wild-type, Fgf3 induction, and Fgfr1 inhibition conditions. **F.** Probing gene expression upon clonal expression of constitutively active Fgfr1 (red, lexop:ca-fgfr1-RFP). Left panel: composite image of pLLP with ca-Fgfr1-expressing clones and *cxcr7b* RNA smFISH on membrane-labeled pLLP. Plot below: corresponding signal intensity profiles along the migration axis. Right panel: activation of *pea3* and *cxcr7b* expression (white, smFISH) through clonally expressed ca-Fgfr1 (red, lexop:ca-fgfr1-RFP) at the leading edge of pLLP (green, cldnb:lyn-gfp). **G.** Illustration of *cxcr7b* expression domain (red cells) upon FGF perturbations (blue-lined cells: *ca-fgfr1* clones, green cells: *fgf3* overexpression). Scale bars: (A, C) 20 µm, (F) 5 µm.

Differential gene expression analysis identified transcripts significantly affected by Fgfr1 inhibition and Fgf3 induction (Table S1, Figure S2A, see Methods). Of the 446 differentially expressed genes shared between both conditions, 329 showed opposing changes after Fgf3 induction versus Fgfr1 inhibition, allowing straightforward classification as either positive or negative FGF targets. For example, expression of the Ets transcription factor *pea3* (*etv4*)^14^, a known positive target, increased with FGF induction and decreased upon Fgfr1 inhibition. By contrast, the transcription factor *lef1* decreased in response to Fgf3 induction and increased with Fgfr1 inhibition, consistent with its previous identification as a negative FGF target^31–33^ (Figure 1B, Figure S2B). We also identified 117 genes that were repressed under both conditions, precluding their classification as either positive or negative targets (Figure 1B, Table S2). Among these was the scavenger receptor *cxcr7b*, previously shown to generate local chemokine gradients that guide the ‘self-directed’ migration of the pLLP^34,35^. Transcription of *cxcr7b* was significantly reduced after Fgfr1 inhibition, consistent with earlier findings that its expression depends on FGF signaling^33^. However, its expression also decreased after Fgf3 induction (Figure 1B-C, Figure S3A-C). This paradoxical response prompted us to examine this novel class of differentially expressed genes more closely, using quantitative approaches comparing *cxcr7b* with the canonical positive target *pea3*. Single-molecule FISH (smFISH) analysis of wild-type pLLPs confirmed that neither gene is normally transcribed at the tissue’s leading domain, and that their expression is turned on in regions where neuromast assembly is promoted by FGF signaling (Figure 1D-E).

Consistently, Fgfr1 inhibition led to a reduction of both genes in these regions. However, while Fgf3 induction strongly increased *pea3* expression (Figure 1D), the same manipulation repressed *cxcr7b* across the tissue^28,29^ (Figure 1E), confirming its unexpected response to changes in FGF activity. The non-canonical nature of this regulation was further underscored by comparison with the closely related paralog *cxcr7a*, which responded to FGF inhibition and induction like a canonical positive target, similar to *pea3* (Figure S3D–G). Since both Fgfr1 inhibition and Fgf3 induction suppressed *cxcr7b* expression, we investigated the impact of directly activating FGF receptor signaling by driving mosaic expression of a constitutively active form of the receptor (ca-Fgfr1a) in the pLLP. As shown in Figure 1F, ca-Fgfr1a triggered cell-autonomous activation of both *pea3* and *cxcr7b* in leading-domain cells, where neither gene is normally expressed. These data support the conclusion that *cxcr7b* transcription is promoted by Fgfr1 signaling but repressed by increased expression of its ligand Fgf3 (Figure 1G).

If *cxcr7b* expression is promoted by Fgfr1 activation, how does Fgf3 induction lead to its downregulation? To address this, we compared *cxcr7b* and Fgf3 expression patterns at cellular resolution using quantitative smFISH together with BAC live reporters (Figure 2A–D, Figure S3H). This analysis revealed that while *cxcr7b* was strongly expressed in neuromast progenitors, similar to *pea3*, its levels were markedly lower in the central cells of rosette-like assemblies (Figure 2A–B, Figure S4A). To more precisely identify ligand-producing cells, we employed a *fgf3:Fgf3-GFP* line, in which a fluorescently tagged gene recapitulates the endogenous expression pattern and rescues ligand function in deficient embryos ^29^ . As expected, secreted Fgf3-GFP accumulated in extracellular luminal spaces, but gene expression was confined to central cells that lacked *cxcr7b* (Figure 2C–D, Figure S4B). A similar mutually exclusive pattern of *cxcr7b* and Fgf3-GFP was observed in other regions of the lateral line (Figure 2A vs 2C, Figure 2E). Despite the absence of *cxcr7b* in rosette centers, smFISH revealed strong expression of *pea3*, indicating that loss of *cxcr7b* was not simply due to reduced autocrine FGF signaling (Figure S4A, Figure 2B).

**Figure 2.**
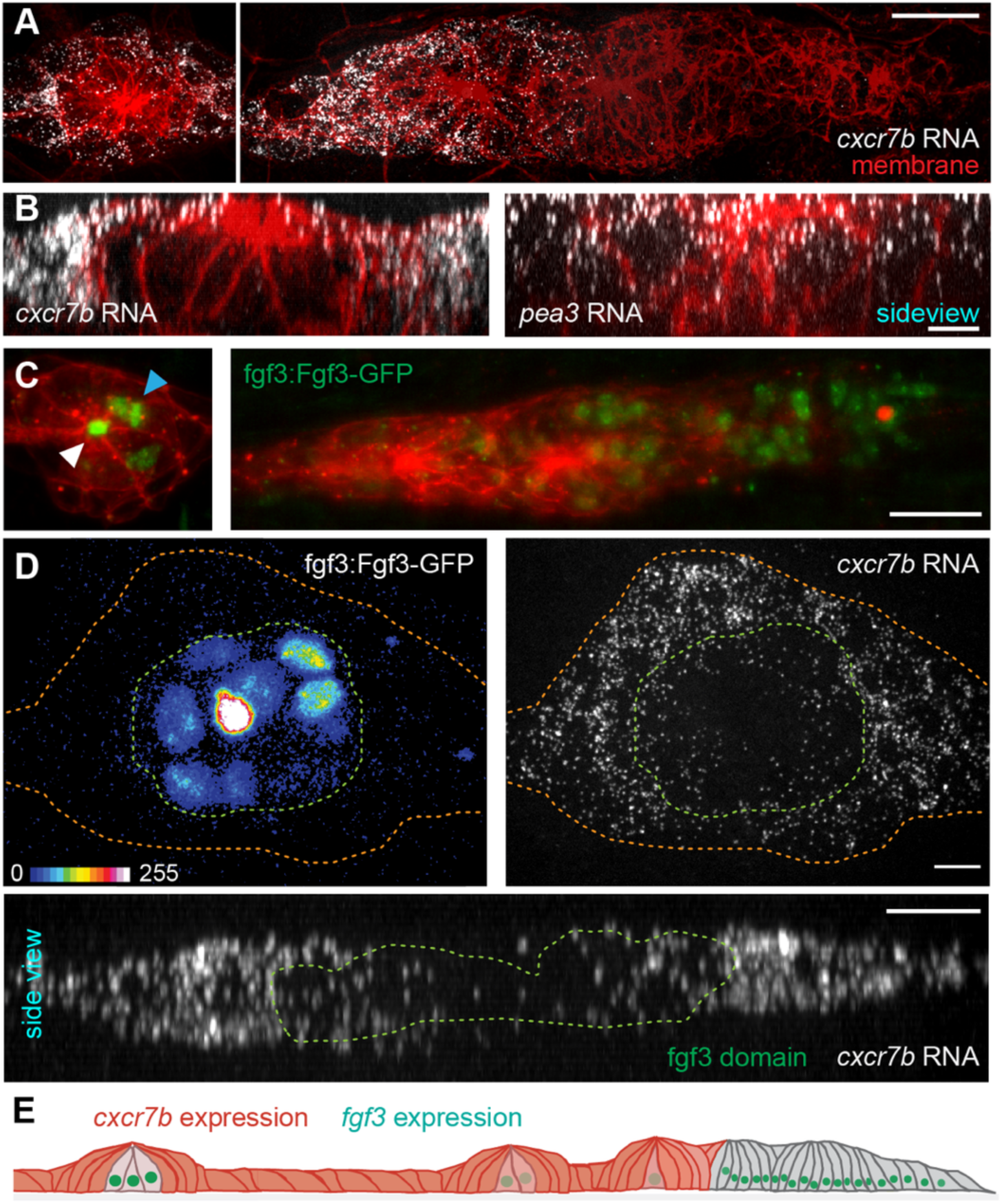
Fgf3 and *cxcr7b* have mutually exclusive expression patterns in the lateral line. **A.** Maximal projection image of *cxcr7b* RNA smFISH (white) in lateral line (left: deposited organ progenitor, right: primordium). **B.** Side view image (middle slice) of *cxcr7b* and *pea3* RNA (white) in deposited organ progenitors (cldnb:lyn-GFP, red), displaying the expression patterns in central versus neighboring cells (see Figure S4A for top view). **C.** fgf3:Fgf3-GFP expression and localization in the lateral line (left: deposited organ progenitor, right: primordium). Blue arrow points to extracellular (luminal) Fgf3 and white arrow to intracellular Fgf3. **D.** Mapping Fgf3 expression domain (green lines) versus *cxcr7b* expression pattern in the same deposited organ (red lines) using fgf3:Fgf3-GFP rescue in Fgf3/Fgf10 knock-down embryos, showing centrally positioned FGF source cells with Fgf3-GFP (displayed in 16 color look-up-table, left panel), and *cxcr7b* RNA smFISH (white, right panel). Top panels: maximal projections; bottom panel: side view, highlighting diminished *cxcr7b* expression in Fgf3 source cells (see also Figure S4B). **E.** Illustration of *cxcr7b* (red) and *fgf3* (green) expression domains in lateral line (grey). Scale bars: (A, C) 20 µm, (B, D) 5 µm.

Beyond its gene expression pattern, imaging Fgf3 ligand revealed a substantial pool of Fgf3-GFP localization to the nuclei of expressing cells, a finding not predicted for a canonical signaling ligand function ^28,31,36^ (Figure 2C–D, Figure 3A). This nuclear localization as well as its mutually exclusive pattern with *cxcr7b* expression was evident also in other embryonic tissues, including the developing pharyngeal arches and otic placode, where nuclear Fgf3-GFP–positive cells lacked *cxcr7b* expression, while juxtaposed neighbors expressed it strongly. These observations suggest that the underlying mechanism operates broadly in *fgf3*-expressing tissues (Figure 3B, Figure S4C). We next asked whether a similar expression pattern could explain the non-canonical regulation of other genes in this class. The T-box transcription factor *tbx1*, like *cxcr7b*, was downregulated both by receptor inhibition and by Fgf3-GFP induction (Figure S4D–E). smFISH confirmed that *tbx1* expression was higher in cells neighboring Fgf3-GFP–positive cells and lower in source cells (Figure 3C–D).

**Figure 3.**
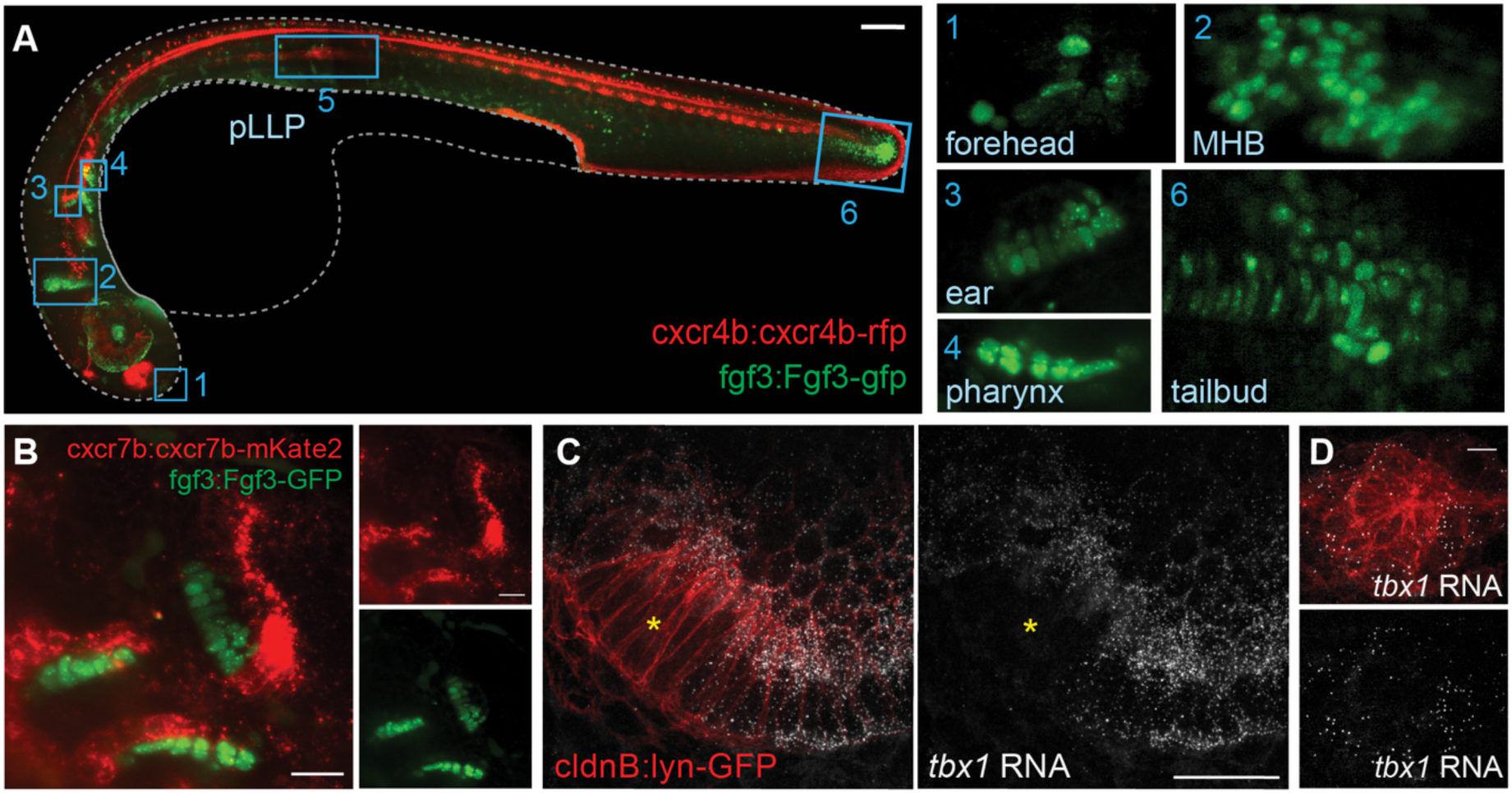
Fgf3-GFP shows nuclear localization and a mutually exclusive pattern with cxcr7b expression in multiple embryonic tissues. **A.** fgf3:Fgf3-GFP 30 hpf zebrafish transgenic line (green) with cxcr4b:cxcrb4-RFP counter-label (red), marking anatomical regions. Known *fgf3* expression domains are revealed by the nuclear localization of Fgf3-GFP in expressing cells. Right panels show the close-up images of the boxed regions (for box #5, see pLLP in Figure 2C). **B.** fgf3:Fgf3-GFP (green), cxcr7b:cxcr7b-mKate2 (red) double transgenic fish showing mutually exclusive neighboring patterns at the developing ear and pharynx (see also Figure S4C). **C, D.** *tbx1* RNA smFISH (white) with cldnb:lyn-GFP membrane label (red) in developing ear and a deposited organ progenitor of lateral line (see Figure S4D). *tbx1* gene expression is detected surrounding Fgf3 expressing regions (ear in Figure 3B, lateral line in Figure 2) but shows a mutually exclusive pattern with the Fgf3-expressing cells (region marked with a yellow star). Scale bars: (A) 100 µm, (B, C) 20 µm, (D) 5 µm.

We next investigated whether this nuclear localization is a feature shared by other FGF family members. We first performed multiple sequence alignments of the FGF family members, which confirmed the previously defined seven subgroups ^37^ (Figure 4A, Figure S5A). We then selected a representative member from each subgroup and predicted their structures ^38^ (Figure S5). This analysis identified features indicative of nuclear localization signals (NLSs), such as positively charged regions on the protein surfaces in an extended conformation ^39^ (Figure S5B-E). Except for Fgf19, which is classified as an “endocrine FGF” due to its reduced affinity for HSPGs and its systemic hormonal activity^40,41^, all analyzed FGF family members displayed putative nuclear-targeting features that overlap with their HSPG interaction surface. To test these predictions, we tagged representative FGFs with GFP between the secretory signal and the globular domain and imaged their subcellular localization after overexpression in early embryos (Figure 4B–C). The subcellular localization of FGF2, which lacks a typical secretory signal, confirmed the reported nuclear targeting of b-FGFs^42^, whereas FGF19 confirmed the exclusive extracellular targeting of the endocrine FGFs. All other FGFs showed varying degrees of nuclear localization, some consistent with previous observations^43,44^. To test the potential context dependence and the impact of the model system on nuclear localization, we imaged mouse FGF3-GFP in mouse embryonic stem cells (mESCs) upon its inducible expression. In mESCs, FGF3 was exclusively targeted to the secretory pathway, with no detectable nuclear localization. Upon differentiation of mESCs through embryoid body formation, however, a subset of cells began to show nuclear localization (Figure S6). These experiments suggest that nuclear localization may be a more widespread latent property of FGFs. FRAP analysis further showed that the nuclear pool of Fgf3-GFP exhibited diffusion kinetics slower than NLS-tagged GFP, yet similar to a widely studied transcription factor, N-Myc, suggesting potential interactions with nuclear components (Figure 4D).

**Figure 4.**
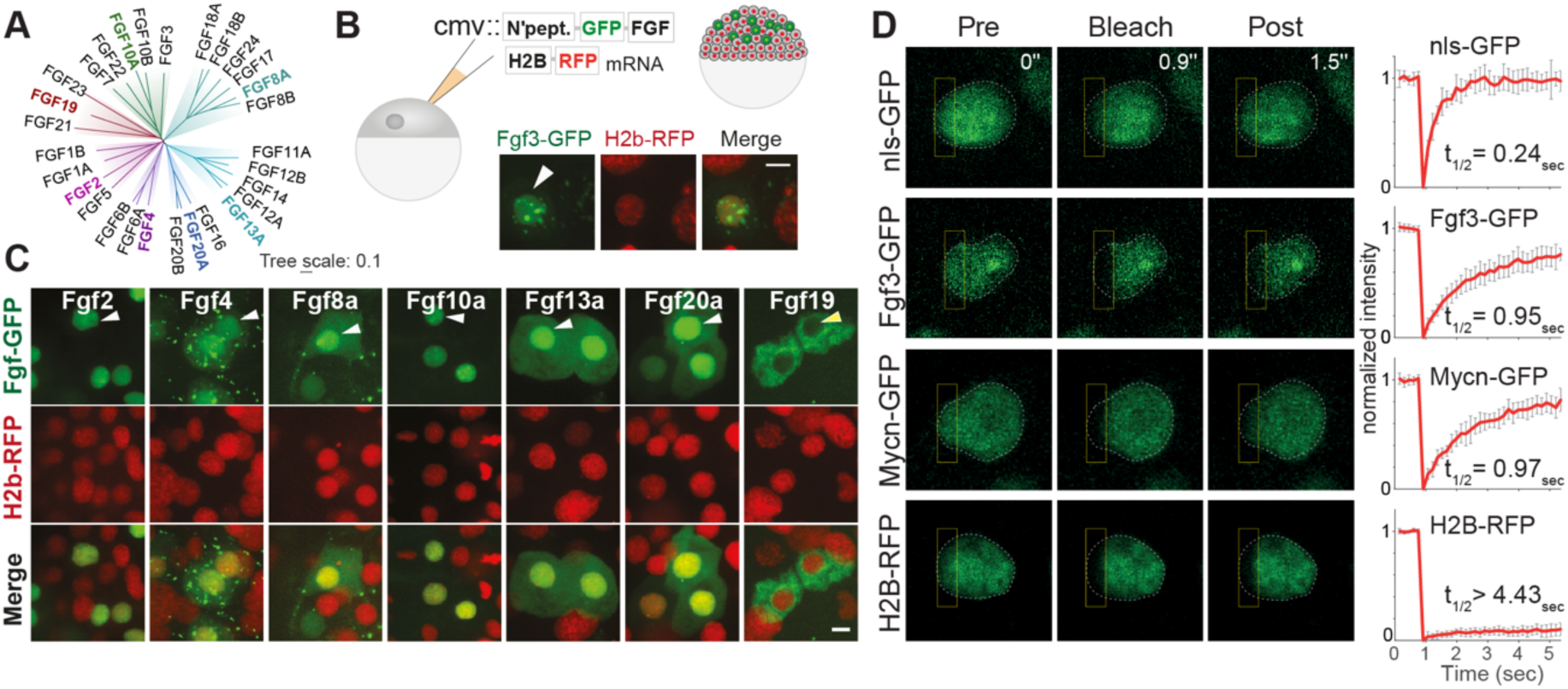
Nuclear localization of Fgf3 has the potential to be a widespread feature among FGFs. **A.** Unrooted phylogenetic tree of FGF family members. Representative members (bold, colored) of each sub-family were chosen for structural modelling and cellular localization experiments. **B.** Scheme summarizing tagging and misexpression of FGF family members: transient misexpression of cmv:fgf-gfp DNA and nuclear counter-label H2b-RFP RNA (red) in 4hpf embryos. The image shows nuclear localization of Fgf3-GFP under conditions of high, transient expression in early embryos. **C.** Cellular localization of GFP-tagged FGF family members (green) with H2b-RFP nuclear marker (red). Arrowheads (white: positive, yellow: negative) indicate nuclear localization of FGFs **D.** Images and quantification of nuclear diffusion using fluorescence recovery after photobleaching (FRAP), contrasting Fgf3, a nuclear tag (nls), Mycn transcription factor and Histone H2B, tagged with a fluorescent protein. Images show pre- and post-bleach time-points. Bleached area is marked with a rectangle. Plots show FRAP of 10 replicates. H2b-RFP did not recover within imaging duration, as expected. Calculated mobile fraction: 0.98, 0.83, 0.86, t_1/2_: 0.24, 0.95, 0.97, respectively. Scale bars: (B, C) 10 µm.

The mutually exclusive expression of *fgf3* with a subset of its signaling targets, combined with nuclear ligand localization in source cells, indicates that Fgf3 may regulate this gene class through a mechanism distinct from canonical extracellular signaling. To probe this directly, we generated additional sources of inducible Fgf3-GFP in nascent neuromasts using cell transplantation (Figure 5A). Donor source cells and neighbor host cells were distinguished by specific reporters, and the impact of Fgf3 induction on *pea3* and *cxcr7b* expression was quantified in individual 3D-segmented cells using smFISH (Figure 5A). Upon induction, *pea3* expression increased in both neighbors and source cells, indicating paracrine and autocrine receptor activation (Figure 5B). Increased expression of the positive target *pea3* in neighbor cells confirmed that paracrine FGF signaling is elevated, consistent with the previous demonstration that neuromast cells can share signals via common extracellular microlumen to ensure coordinated signaling^29^. An equivalent increase in *pea3* mRNA level in source cells suggests that these cells are just as responsive to the ligand they secrete as neighboring cells (Figure 5B). By contrast, *cxcr7b* expression was significantly reduced only in Fgf3-GFP source cells, not in neighbors (Figure 5B, Figure S7). These results support a dual mode of Fgf3 action: activating target genes in receiving cells via receptor-mediated autocrine and paracrine signaling, while cell-autonomously repressing the targets in ligand-producing source cells.

**Figure 5.**
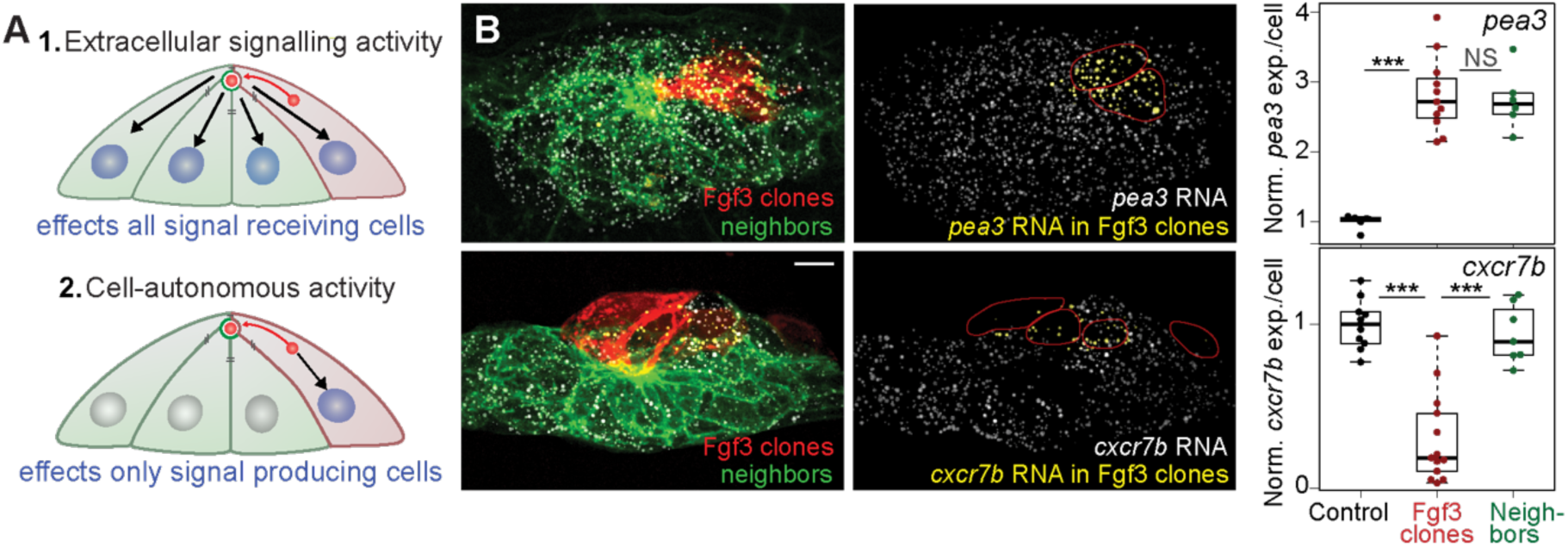
Clonal misexpression of Fgf3 leads to increased signaling in all organ progenitor cells but cell autonomous downregulation of *cxcr7b* expression specifically in source cells. **A.** Schema showing possible outcomes of ectopic Fgf3 expression in clones. Fgf3-misexpressing cells are drawn in red, other cells in the same organ progenitor (neighbors) are drawn in green. **B.** *pea3* and *cxcr7b* RNA smFISH in deposited mosaic organ progenitors with Fgf3-induced (Fgf3-GFP/lyn-RFP/nls-tdTomato) cell clones (red) among cldnb:lyn-GFP labeled neighbor cells (green). RNA spots identified by 3D segmentation are shown in yellow if located within FGF-expressing clones, or white if in surrounding cells of the same organ progenitor. Red outlines (right panels) mark the positions of the Fgf3-expressing clones. Box plots show the quantification of *pea3* and *cxcr7b* levels per cell in deposited organ progenitors contrasting FGF expressing clones with the rest of the neighboring cells in the same organ progenitor (*n* = 5, 11, 6; *n* = 10, 13, 7; Wilcoxon test, *\*\*\*P <* 0.001). Scale bar: 5 µm.

The data described above demonstrate that Fgf3 localizes to the nucleus in expressing cells and can repress specific target genes in a cell-autonomous manner. To probe the causal link between these findings, we performed experiments manipulating the intracellular transport route of Fgf3. High-resolution imaging revealed that full-length Fgf3-GFP localized both to the nucleus and to the secretory pathway in source cells, with the secreted pool concentrated in the central microlumen^45^ (Figure 6A, Figure S8A). To alter ligand routing, we manipulated its N-terminal signal peptide, which normally directs secretion and is cleaved from the ligand upon endoplasmic reticulum entry. Fusion of the signal peptide to GFP (secGFP) labeled only the secretory pathway and microlumen, with no detectable nuclear localization (Figure 6B, Figure S8B). Conversely, deleting the signal peptide from full-length Fgf3-GFP (nonsec–Fgf3-GFP) abolished secretion with exclusive nuclear localization and no detectable labelling of the secretory compartments or microlumina (Figure 6C, Figure S8C). This contrast is consistent with nuclear localization acting as the default transport pathway when secretion is diminished.

**Figure 6.**
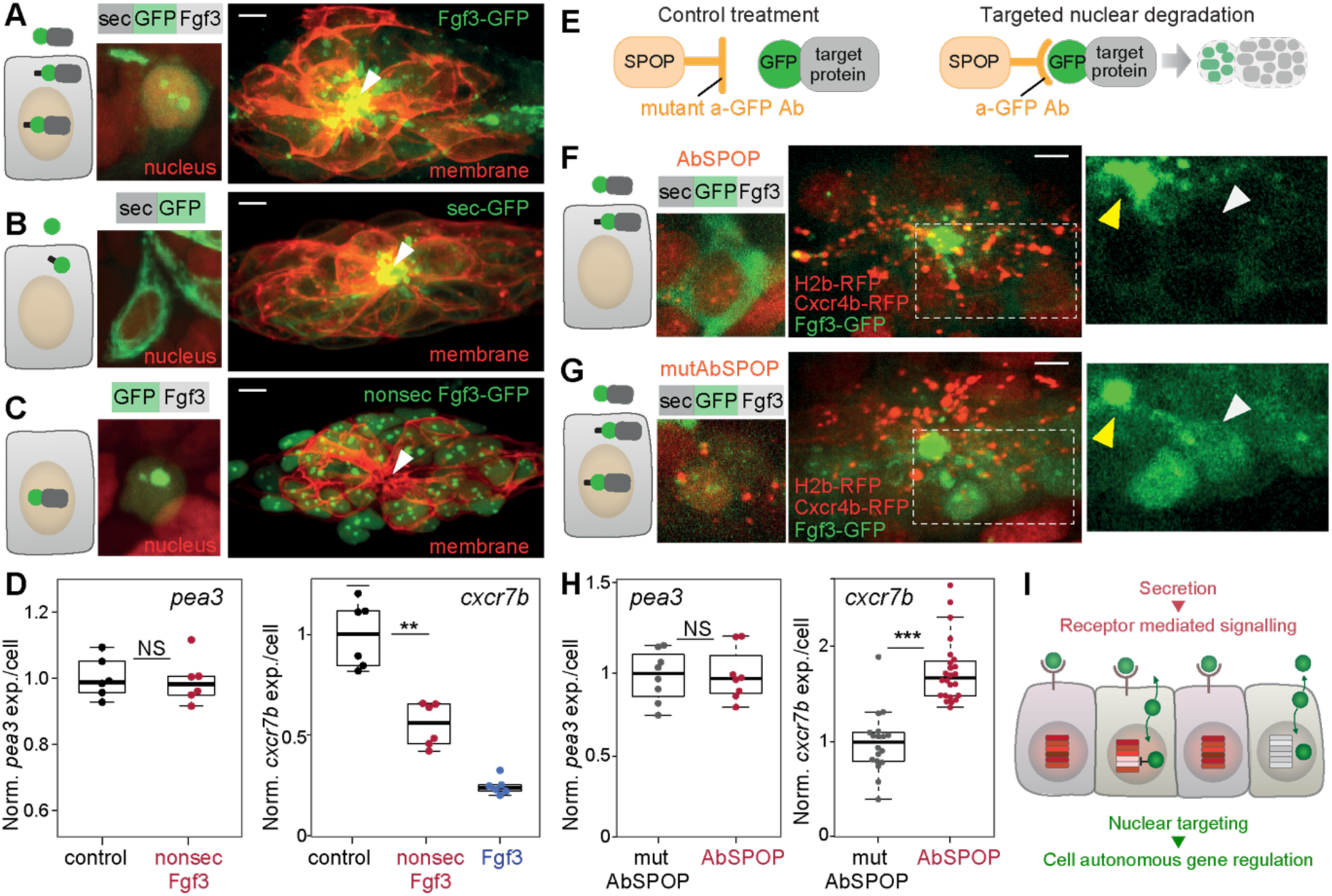
Nuclear localization of Fgf3 in source cells exerts a distinct impact on gene expression compared with its signaling function. **A, B, C.** Localization of GFP-tagged Fgf3 constructs. Fgf3-GFP (sec-GFP-Fgf3) represents full-length Fgf3, tagged with GFP between the signal peptide and the globular domain. sec-GFP is GFP-tagged signal peptide and nonsec-Fgf3-GFP (GFP-Fgf3) is GFP-tagged globular domain of Fgf3 lacking the signal peptide. Left panel: schematic of cellular localization. Middle: cellular imaging of tagged proteins (green) with nuclear counter-label (nls-tdTomato, red). Right: multicellular imaging of the GFP-tagged proteins (green) in assembled organs with membrane counter-label (lyn-RFP). Arrowheads: extracellular space in rosette centers (microlumen). **D**. Boxplots showing quantification of *pea3* expression (left) upon nonsec-Fgf3-GFP expression in organ progenitors and *cxcr7b* expression (right) upon nonsec-Fgf3-GFP and Fgf3-GFP expression (*n* = 6 each). **E.** Scheme of anti-GFP nanobody targeted nuclear degradation. Nuclear-localized proteins were degraded upon recognition of GFP tag via AbSPOP. Mutant anti-GFP nanobody-SPOP fusion (mutAbSPOP) served as non-targeting control. **F, G.** Fgf3-GFP localization upon lexop:fgf3-gfp expression with targeted nuclear degradation (F) versus non-targeting (G). Left: schematic of cellular localizations and imaging of tagged proteins (green) with nuclear counter-label (nls-tdTomato, red). Right: multicellular imaging of the Fgf3-GFP (green) in organ progenitors with counter-labels (H2B-RFP and cxcr4b-RFP) and close-up views of the boxed regions highlighting Fgf3-GFP cellular localization. Microluminal (extracellular space) and nuclear spaces are indicated by yellow and white arrows, respectively. H, *pea3* and *cxcr7b* expression in control versus nuclear degraded Fgf3-GFP expressing organ progenitors (*n* = 8 each; *n* = 18 and 24, respectively)**. I.** Schematic of the FGF-driven gene regulation model. Scale bars: (A-C, F-G) 5 µm. Statistics: Wilcoxon test, *\*\*P <* 0.01,*\*\*\*P <* 0.001.

Functionally, nonsec–Fgf3-GFP was deficient in inducing receptor-mediated signaling as evidenced by the absence of detectable increase in *pea3* mRNA in expressing neuromasts (Figure 6D). This absence rules out the possibility of Fgf3 signaling via non-canonical secretion pathways. The nonsec-Fgf3-GFP protein, however, repressed *cxcr7b* expression (Figure 6D), suggesting that nuclear Fgf3 can mediate repression of specific target genes in signal source cells.

Performing the complementary experiment, preventing nuclear targeting specifically while leaving signaling activity intact, proved more challenging. Unlike the discrete signal peptide, the residues for the nuclear localization are distributed across the protein and overlap with other structural features (Figure S7). An alternative approach, attaching a nuclear export sequence to Fgf3-GFP, reduced nuclear localization as anticipated but also hindered this variant from entering the secretory pathway, precluding functional distinction (Figure S8D). We therefore took an alternative approach to testing the requirement for Fgf3 nuclear localization by selectively depleting the nuclear pool using a nanobody-based degradation strategy. This targeted degradation is based on DeGradFP method, where anti-GFP nanobodies fused to the SPOP ubiquitin ligase (AbSPOP) specifically target GFP-tagged proteins in the nucleus for degradation without affecting cytosolic protein levels (Figure 6E, Figure S9) ^46^. Indeed, expressing AbSPOP in the Fgf3-GFP-induced cells depleted nuclear Fgf3-GFP signal while leaving the secreted luminal pool unaffected (Figure 6F). Validating the specificity of this approach, expressing a nanobody mutant version (mutAbSPOP), which retains functional SPOP but lacks the ability to bind GFP-tagged targets, did not interfere with Fgf3-GFP distribution (Figure 6G). Importantly, selective depletion of nuclear Fgf3-GFP by AbSPOP did not alter *pea3* expression levels, confirming intact extracellular signaling, but increased *cxcr7b* levels as compared to mutAbSPOP (Figure 6H). Together, this reveals Fgf3’s ability to repress the expression of specific target genes in source cells through nuclear localization, a mechanism that is decoupled from its function in extracellular signaling (Figure 6I).

## Discussion

Previous studies on cell signaling have primarily focused on how gene expression patterns are shaped by secreted ligands via paracrine and autocrine signaling^3,36^. By combining live imaging of the Fgf3 ligand with gene expression profiling of its downstream targets, we uncover an underexplored role for this classical paracrine signal in regulating gene expression within a multicellular tissue context. We propose a patterning model in which extracellular FGF regulates the expression of canonical target genes in all signal-receiving cells, while nuclear FGF selectively represses a subset of these targets in ligand-sending source cells (Figure 6I). By innately distinguishing source cells from their neighbors, this mechanism offers a general strategy for establishing self-organized gene expression differences across diverse tissues, such as the ear^47^ and pharynx^47,48^. The dual function of FGF signaling revealed here appears to introduce an additional layer of regulatory control in developmental signaling.

To address the basis of this dual function, we manipulated the subcellular localization of Fgf3, allowing us to deconvolve its nuclear and extracellular roles. Fgf3 secretion was specifically prevented by deleting its defined N-terminal signal peptide. This non-secreted variant localized to the nucleus and repressed *cxcr7b* transcription, consistent with the conclusion that nuclear-localized pool does not enter the secretory pathway. Interestingly, the non-secreted variant appears less effective at suppressing *cxcr7b* transcription than the full-length Fgf3-GFP (Figure 6D), raising the possibility that the signal sequence, which is normally cleaved from the secreted ligand only upon translocation into the ER, may enhance the activity of the nuclear-localized form. The converse rerouting experiment, preventing nuclear targeting, was challenging, nevertheless the targeted protein degradation approach directly demonstrated that nuclear Fgf3 is required to suppress cxcr7b expression, while leaving paracrine signaling intact. Looking ahead, it will be important to determine whether developmental regulation of Fgf3 partitioning between nuclear and secretory pathways provides adaptive control. Although beyond the scope of this study, future work defining the interaction of FGFs with chromatin and nuclear partners will clarify how nuclear-localized FGFs regulate specific target genes.

The cell-autonomous nuclear role of Fgf3-GFP, both at physiological expression levels and upon targeted misexpression, prompted us to investigate the other FGF family members. Guided by the structural predictions, misexpression experiments in early embryos suggests that nuclear functions could represent a broader property of FGF ligands. Future studies leveraging endogenous detection of FGF proteins across cell-types will be required to test whether this nuclear localization occurs under physiological conditions and in specific cellular contexts.

Together, these findings highlight an intriguing and underexplored dimension of FGF function in development with potential implications for disease. The detection of nuclear-localized FGF ligands prompts a re-evaluation of the roles and subcellular localization of other signaling molecules, particularly those containing exposed positively charged regions that could promote nuclear targeting, a sequence feature shared by many ECM-associated growth factors and signaling molecules^49^.

Revisiting genetic disorders linked to mutations in such molecules could yield new insights^50^, especially considering the potential cell-autonomous activities distinct from their conventional paracrine functions^51^.

## Data availability

Datasets generated in this study were deposited at Gene Expression Omnibus (GEO, https://www.ncbi.nlm.nih.gov/geo/) under accession no GSE207538.

## Contributions

S.D. and D.G. designed the study. S.D. performed all experiments, assisted by A.R.M for colorimetric in situ hybridizations. M.I. and S.D. analyzed the data and assembled the figures with input from P.B. and D.S.. E.G. developed the inducible ca-Fgfr1 system. A.K. generated GFP-tagged FGF ligand plasmids and assisted with 3D organ segmentation. E.K. performed FGF protein sequence and structure comparisons. E.D. cloned *cxcr7a* cDNA. S.D. and D.G. interpreted the data and wrote the paper with input from all authors.

## Competing interests

The authors declare no competing financial interests.

## Acknowledgements

We acknowledge Gilmour laboratory for discussions and Noa Gil and Sena A. Uslu for feedback on the manuscript. We thank Andrea Gruia, Celine Revenu, Ulrike Schulze, E.D., A.K. and M.I. for the help with preprocessing embryos. We thank EMBL core facilities, Vladimir Benes and Jelena Pistolic (Genecore), Alex Perez Gonzalez (FACS), Yury Belyaev, Kota Miura, Stefan Terjung (ALMF). We thank Andrea Gruia (EMBL) and Cornelia Henkel (MLS-UZH) for fish care. S.D. acknowledges the EMBO Long-Term fellowship (ALTF 1101) and the Christiane-Nüsslein-Volhard stipend. D.S. acknowledges EU Horizon 2020 ERC Advanced Grant (884664) and support from Novartis Research Foundation. D.G. acknowledges the SNSF Grant (31003A_176235) and support from University of Zürich. E.K. acknowledges support from Izmir Biomedicine and Genome Center. E.D. acknowledges support from CNR Neuroscience Institute.

## Supplementary Figures

**Figure S1.**
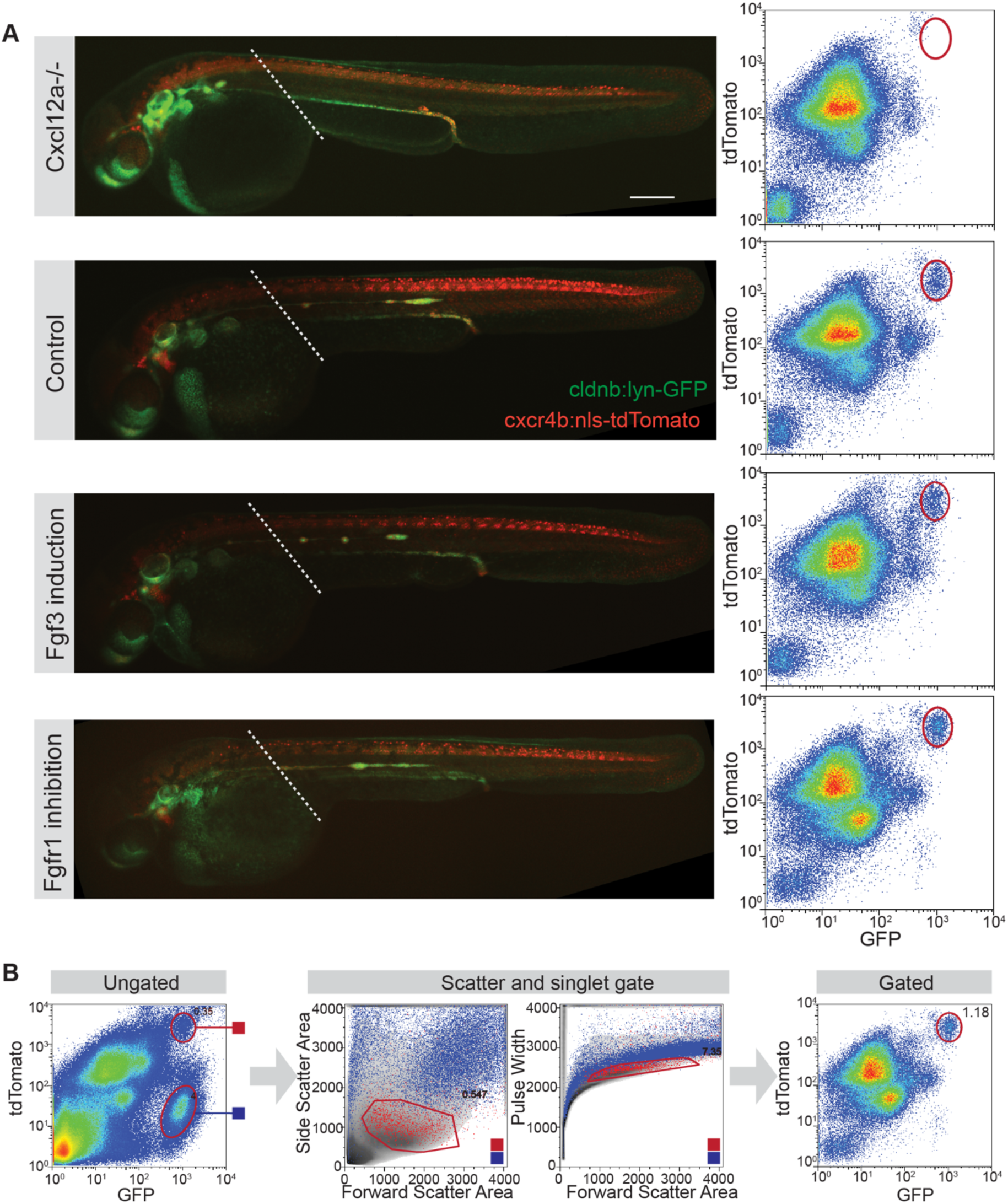
Identification and isolation of the lateral line cells with FACS. **A.** Left: Whole-embryo images of cxcl12a (sdf1a) mutant, wild-type, Fgf3-induced and Fgfr1-inhibited transgenic cldnb:lyn-GFP, cxcr4b:nlsTomato embryos at 30 hpf. Dashed lines indicate the dissection site for trunk isolation. Right: Corresponding fluorescence density plots (blue to red: low to high cell count density) showing gated cells dissociated from trunks. The sorted pLLP population is circled, absent in cxcl12a mutants due to impaired primordium migration. **B.** Sorting gates used to isolate lateral line primordium cells from embryo trunks. Ungated: all trunk cells plotted by tdTomato and GFP fluorescence. The regions corresponding to the reference population (blue box) and the primordium population (red box) are outlined. Scatter and singlet gate: Scatter and singlet gate: scatter plots of analyzed cells (density in grey) with primordium (red dots) and reference population (blue dots) overlaid. Gates used to sort pLLP are outlined with red lines. Gated: fluorescence plot of remaining cells after scatter and singlet gates with sorted primordium population outlined in red. Scale bar: (A) 200 µm.

**Figure S2.**
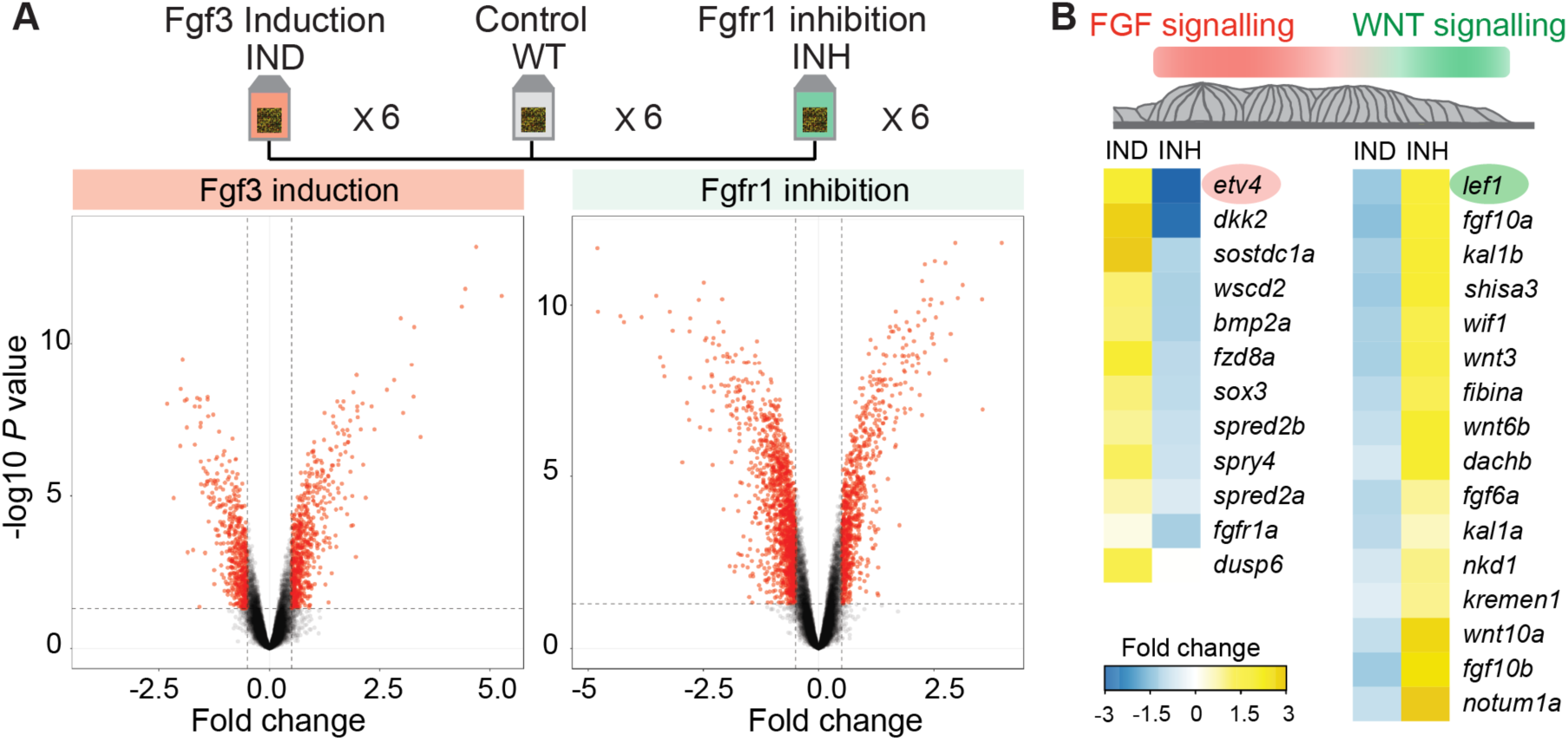
Fgf3 misexpression and Fgfr1 inhibition lead to differential gene regulation in pLLP, impacting known FGF and WNT signaling targets. **A.** Gene expression profiling of Fgf3 induction, wild-type, and Fgfr1 inhibition across six biological replicates. Volcano plots show thresholds used to identify genes differentially expressed under treatment relative to control. **B.** Differential regulation of FGF (red) and WNT (green) signaling components in lateral line primordium. Heatmaps show fold changes relative to wild-type (blue: down-regulation, yellow: up-regulation).

**Figure S3.**
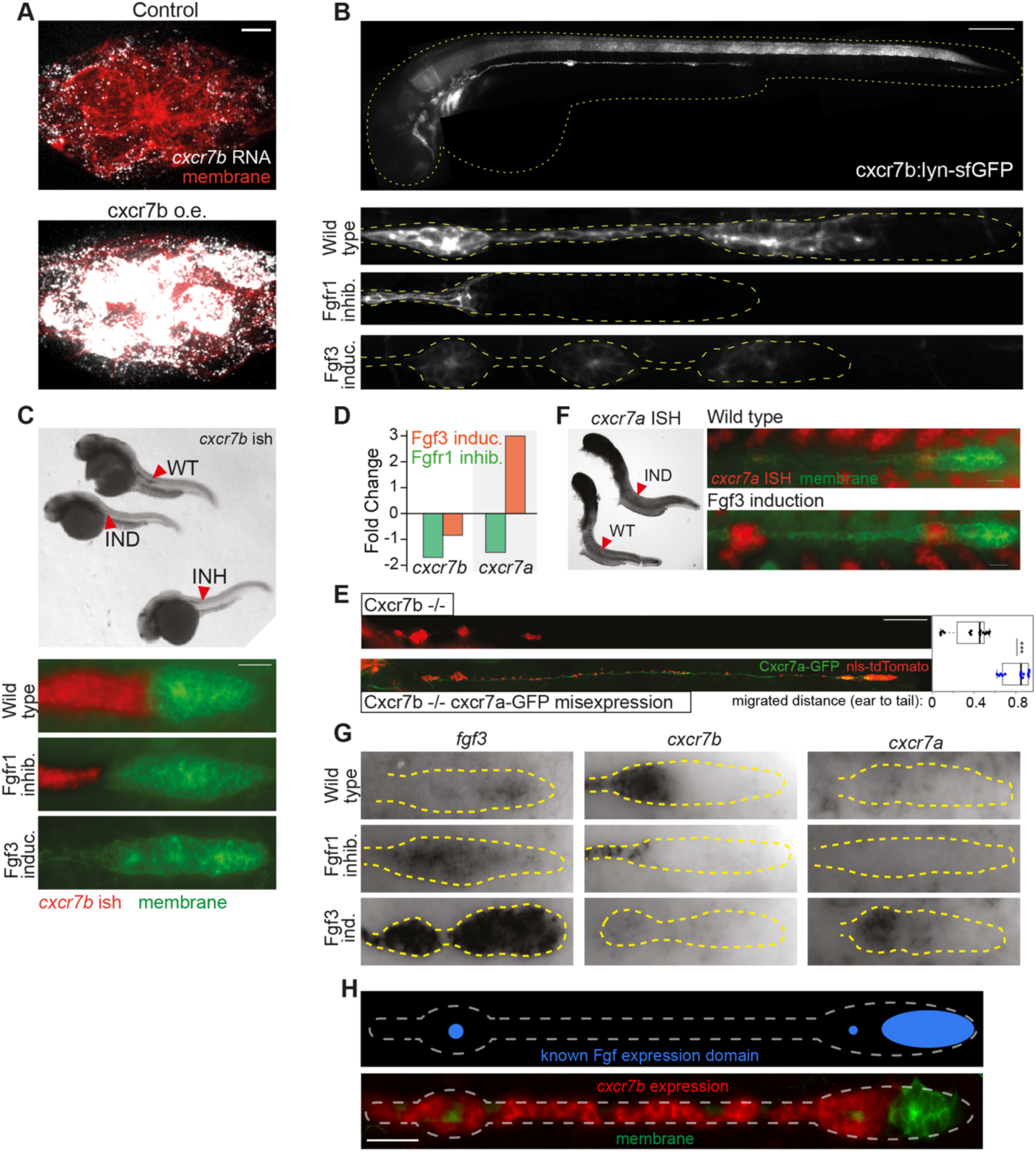
Impact of FGF manipulations on *cxcr7b* expression. **A.** Establishing *cxcr7b* smFISH (grey) using control (cxcr4b:lexPR) and *cxcr7b* overexpressing (cxcr4b:lexPR/lexOP:cxcr7b) lateral line organs (cldnb:lyn-GFP, red), confirming the specific recognition of *cxcr7b* mRNA by the smFISH probes. **B.** Whole embryo image of cxcr7b:lyn-sfGFP (white) transgenic line marking *cxcr7b* expression domains with membrane-GFP (thus a low turn-over rate of the reporter). Lower panels: close-up images of lateral line in cxcr7b:lyn-sfGFP embryos, showing responses to early Fgf3 induction and inhibition. **C.** *cxcr7b* colorimetric RNA in situ hybridization (ISH) of whole embryos (black) and close-ups (red) upon FGF manipulations (cldnb:lyn-GFP, green). **D.** Bar plots of gene expression change of the paralogs, cxcr7b and cxcr7a, upon FGF induction and Fgfr1 inhibition, measured by microarrays. **E.** Rescue of Cxcr7b loss-of-function by Cxcr7a in cell migration. Upper: 2-day-old Cxcr7b mutant embryo with impaired primordium migration. Lower: transient Cxcr7a-GFP expression (gal4/uas:cxcr7a-GFP) rescuing the primordium migration. Migration quantified (*n* = 15 control, 14 rescue). **F.** Left: *cxcr7a* colorimetric ISH (black) on wild-type and Fgf3 induced embryos, showing increased *cxcr7a* expression at lateral line organ progenitors upon Fgf3 induction. Right: close-up images of the lateral line with *cxcr7a* ISH (red) and cldnb:lyn-GFP counter-label (green). **G.** *fgf3*, *cxcr7b* and *cxcr7a* ISH (black) on wild-type, Fgfr1 inhibited and Fgf3 induced lateral line primordium, highlighting the opposite responses of *cxcr7b* and *cxcr7a* to FGF induction when pLLP cells are induced to express the Fgf3 ligand. **H.** The known FGF ligand expression pattern (schema in blue) in contrast to the opposing expression pattern of *cxcr7b* (ISH in red) in the zebrafish lateral line (cldnb:lyn-GFP, green). Scale bars: (A) 5 µm, (B, E) 200 µm, (C) 20 µm, (H) 50 µm. Statistics: Wilcoxon test, *\*\*\*P <* 0.001.

**Figure S4.**
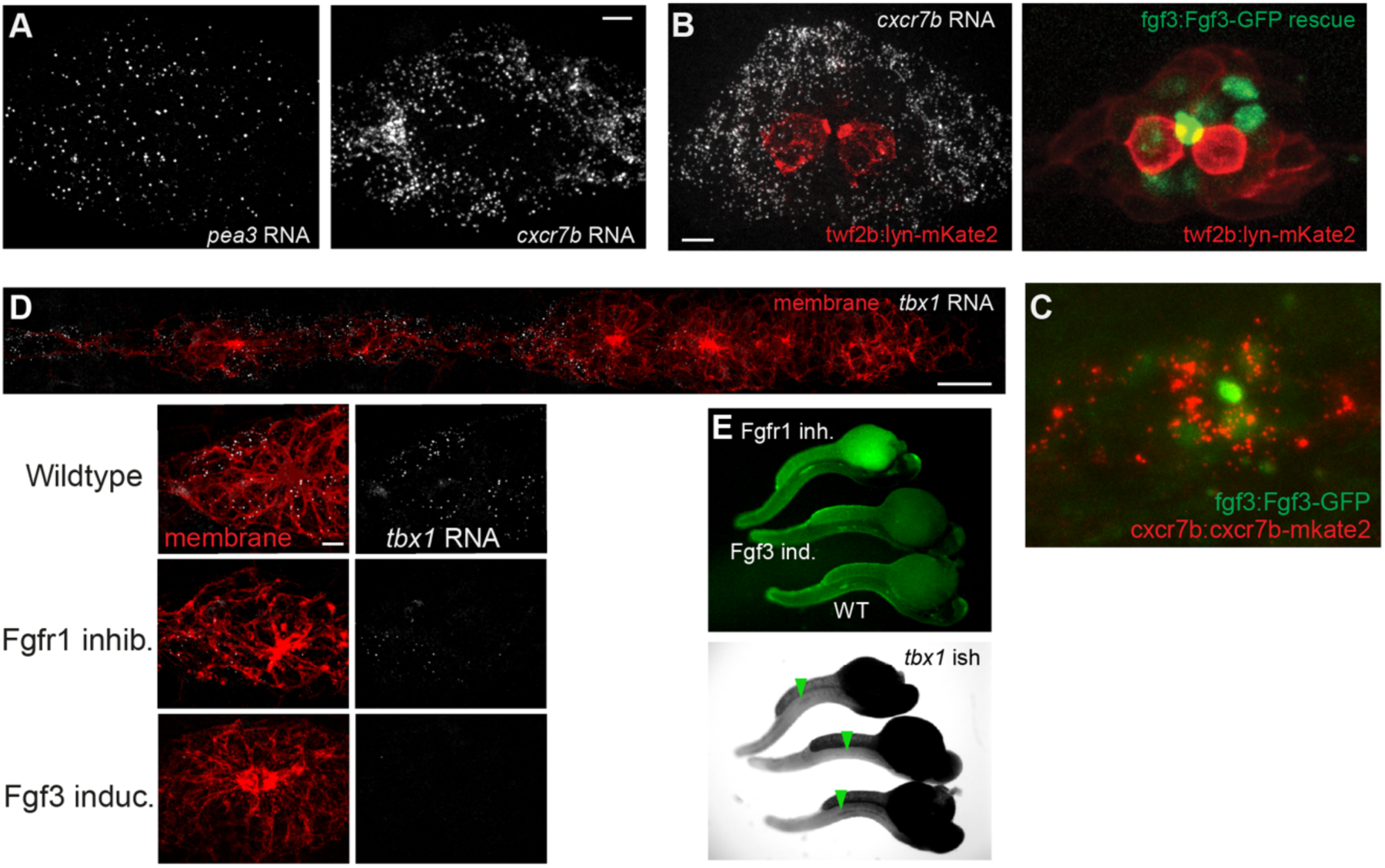
Probing cxcr7b and tbx1 gene expression patterns in comparison to Fgf3. **A.** Single smFISH channels of Figure 2B. **B.** Composite images of *cxcr7b* smFISH (white) and fgf3:Fgf3-GFP rescue (green) from Figure 2D, overlaid with membrane marker of twf2b:lyn-mKate2 (red) to facilitate 3D alignment of two imaging sets from the same organ progenitor. **C.** Composite image of a lateral line organ progenitor marked with cxcr7b:Cxcr7b-mkate2 (red) *and* fgf3:Fgf3-GFP (green). The cellular *cxcr7b* gene expression pattern cannot be resolved using the endogenously tagged Cxcr7b protein in pLLP due to high protein turnover (seen as endocytic punctate signal), unlike the plasma membrane signal in other tissues (Figure 3B). **D.** *tbx1* RNA smFISH (white) on the lateral line (red, cldnb:lyn-GFP) and its response to FGF manipulations at the trailing edge of the primordium, resembling the pattern of *cxcr7b*. **E**. Colorimetric ISH (black) of *tbx1* in whole fish embryos (cldnb:lyn-GFP, green) after FGF manipulations. Arrows point to the pLLP. Scale bars: (A, B, D-lower panel) 5 µm, (D-upper panel) 20 µm.

**Figure S5.**
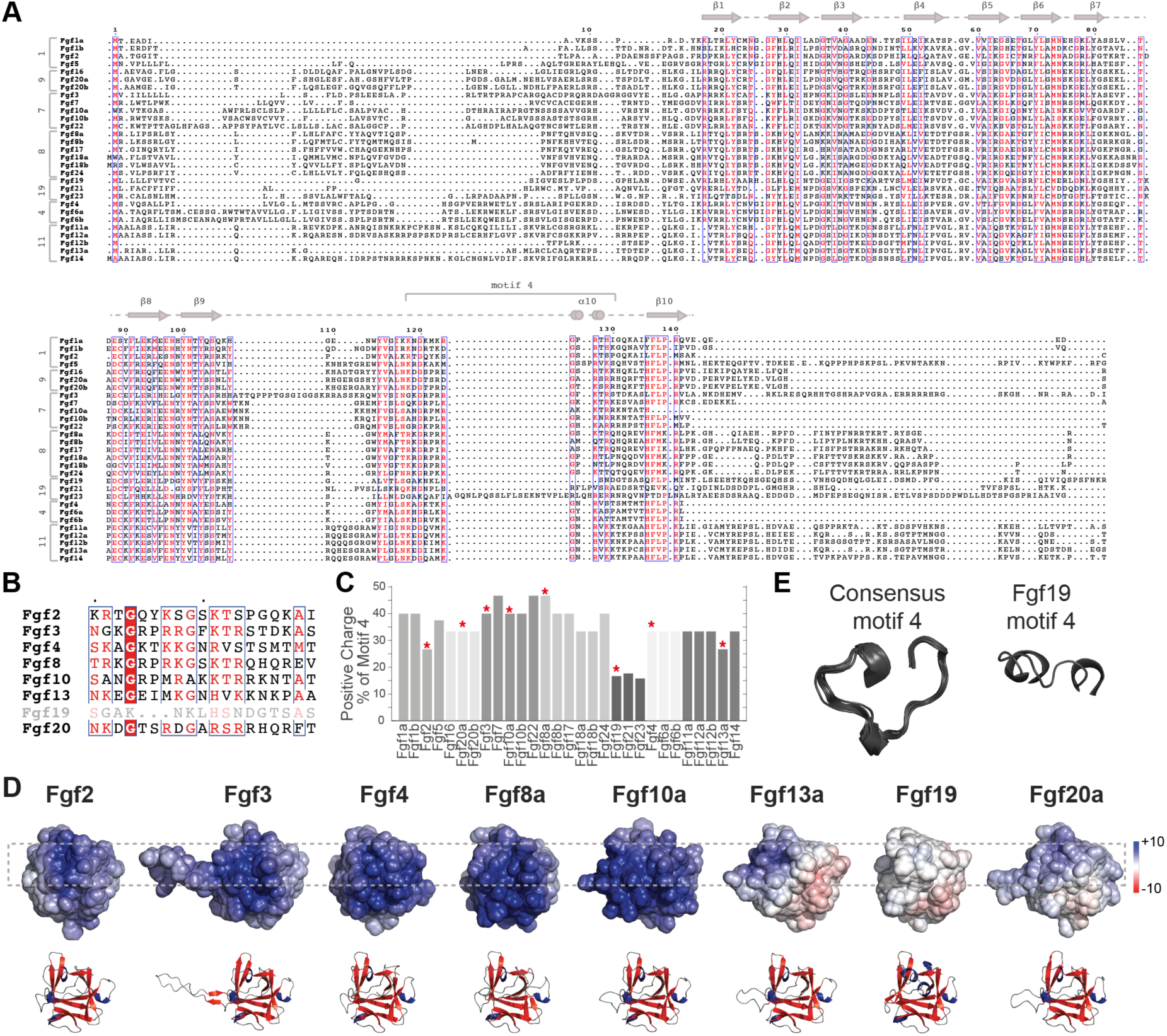
Comparative sequence and structural analysis of zebrafish FGFs. **A.** Multiple sequence alignments of zebrafish FGFs performed using the T-Coffee server^52^ and visualized with ESPript 3.0^53^. Scores above SimilarityGlobalscore are shown in red, and strictly conserved residues in white on red background. A previously predicted motif-4^39^ is highlighted for its potential to act as a nuclear localization signal (positively charged, extended-exposed conformation, conserved). **B.** A closer look at the sequence conservation of motif-4 among the experimentally tested FGFs. **C**. Positive charge percentage of motif-4 among FGFs. FGFs tested experimentally are marked with a star. **D.** Electrostatic surface maps of zebrafish FGFs, with structures predicted by AlphaFold3.0. Surface charge scale spans from -10 (red) to +10 (blue). The region corresponding to motif-4 is boxed, which shows a highly positively charged domain on the surface of the FGFs except Fgf19. **E.** Closer inspection of the motif further showed that its 3D structure was conserved except for Fgf19.

**Figure S6.**
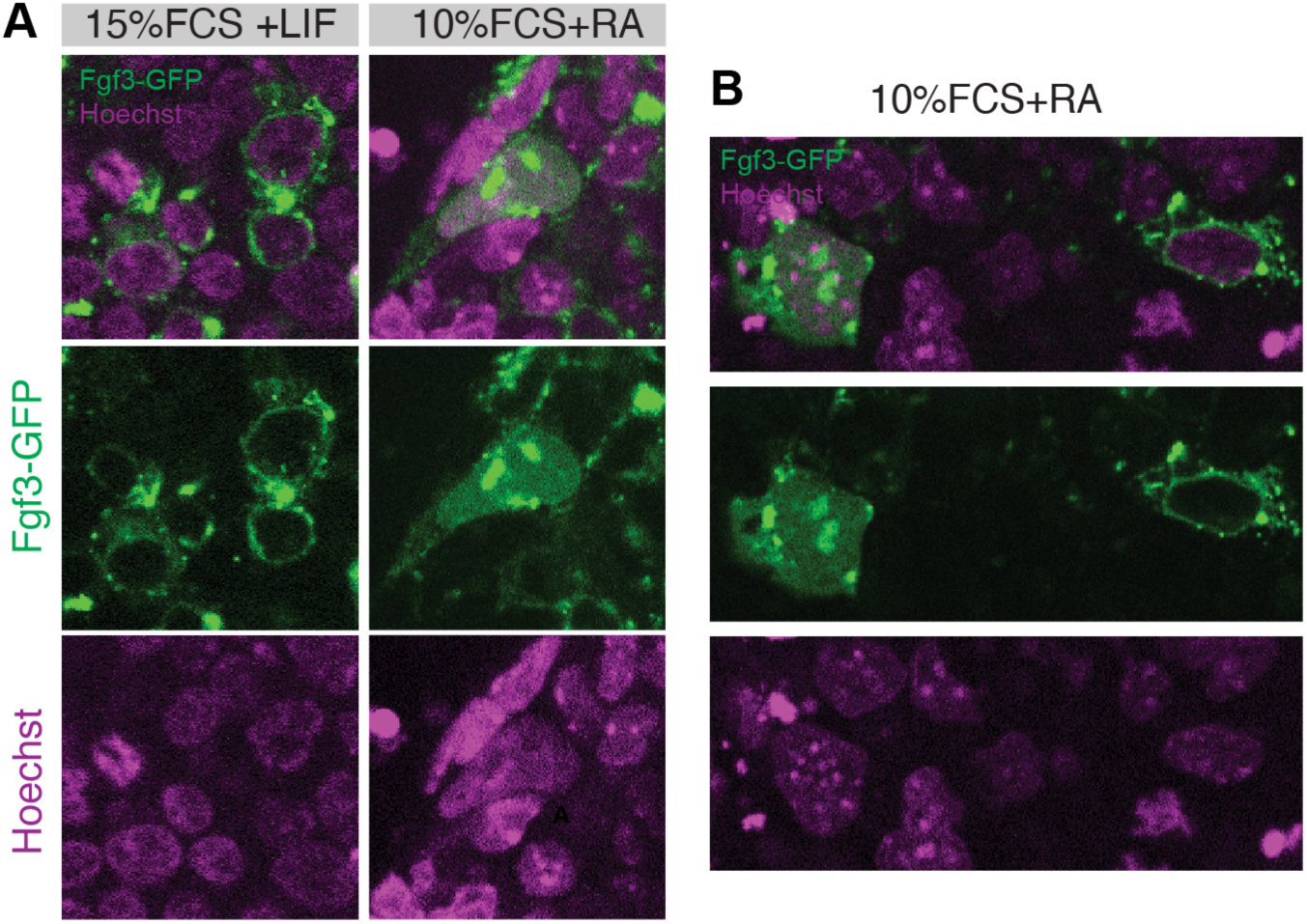
Subcellular localization of mouse FGF3-GFP in mouse embryonic stem cells (mESCs). **A.** Subcellular localization of mouse FGF3-GFP (upon 6 hour doxycycline induced expression) in mESCs cultured in pluripotent maintenance media (15%FCS+LIF) versus spontaneous differentiation induction media (10%FCS+retinoic acid). Nuclei counter-labeled with Hoechst staining (purple). **B.** Cross-sectional image of mESCs in spontaneous differentiation induction media, showing two cells at the same imaging plane, expressing FGF3-GFP, yet with variable subcellular localization.

**Figure S7.**
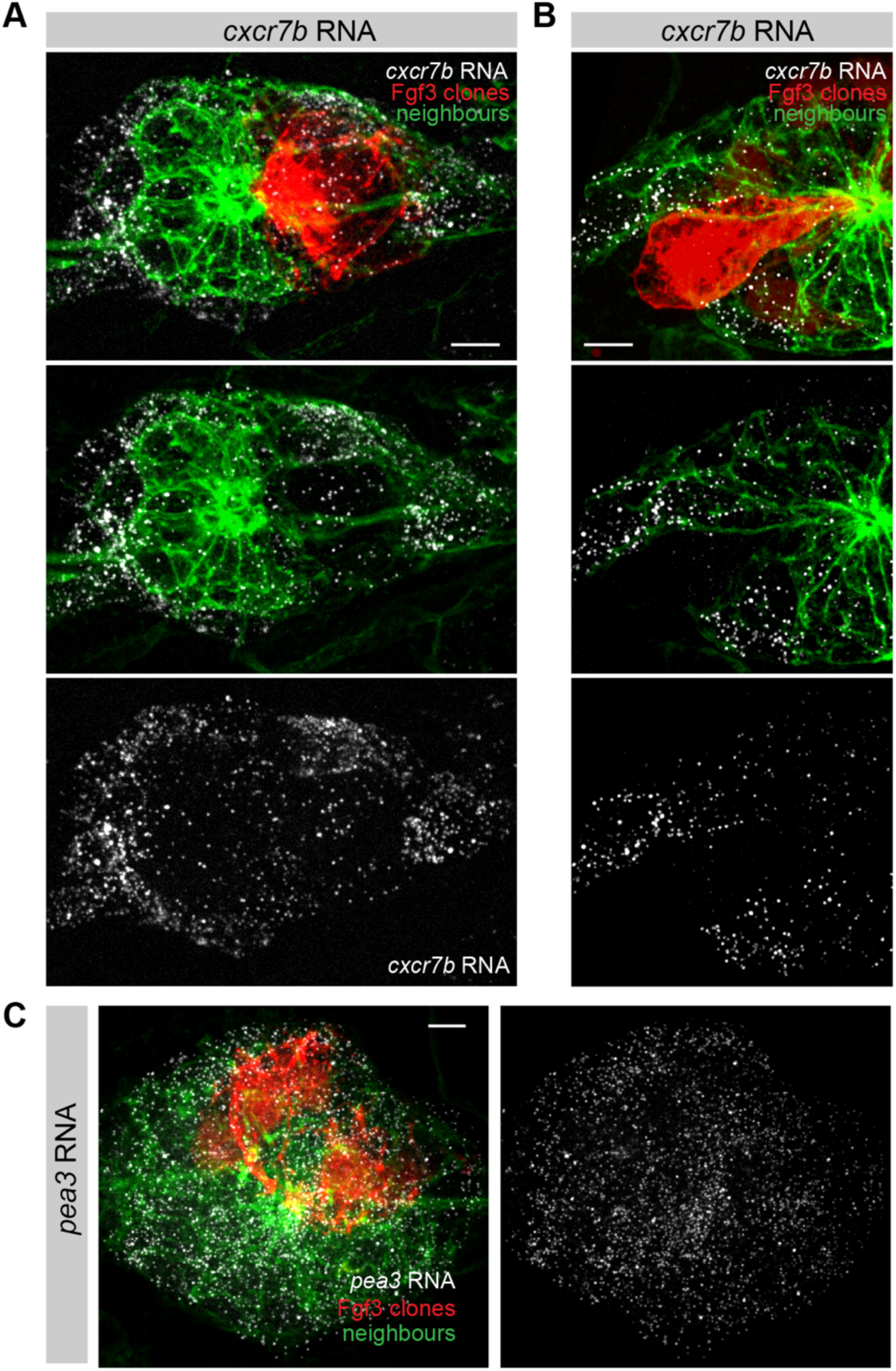
Clonal Fgf3 misexpression demonstrates the cell autonomous repression of *cxcr7b*. **A, B.** Maximum Z-projections of *cxcr7b* smFISH (white) following clonal Fgf3 misexpression (red) and neighboring cells (green) in a mature organ progenitor (A) and at the trailing end of a primordium (B). **C.** *pea3* smFISH (white) on Fgf3 misexpression clones (red) along with surrounding cells (green) in a mature organ progenitor. Scale bars: (A-C) 5 µm.

**Figure S8.**
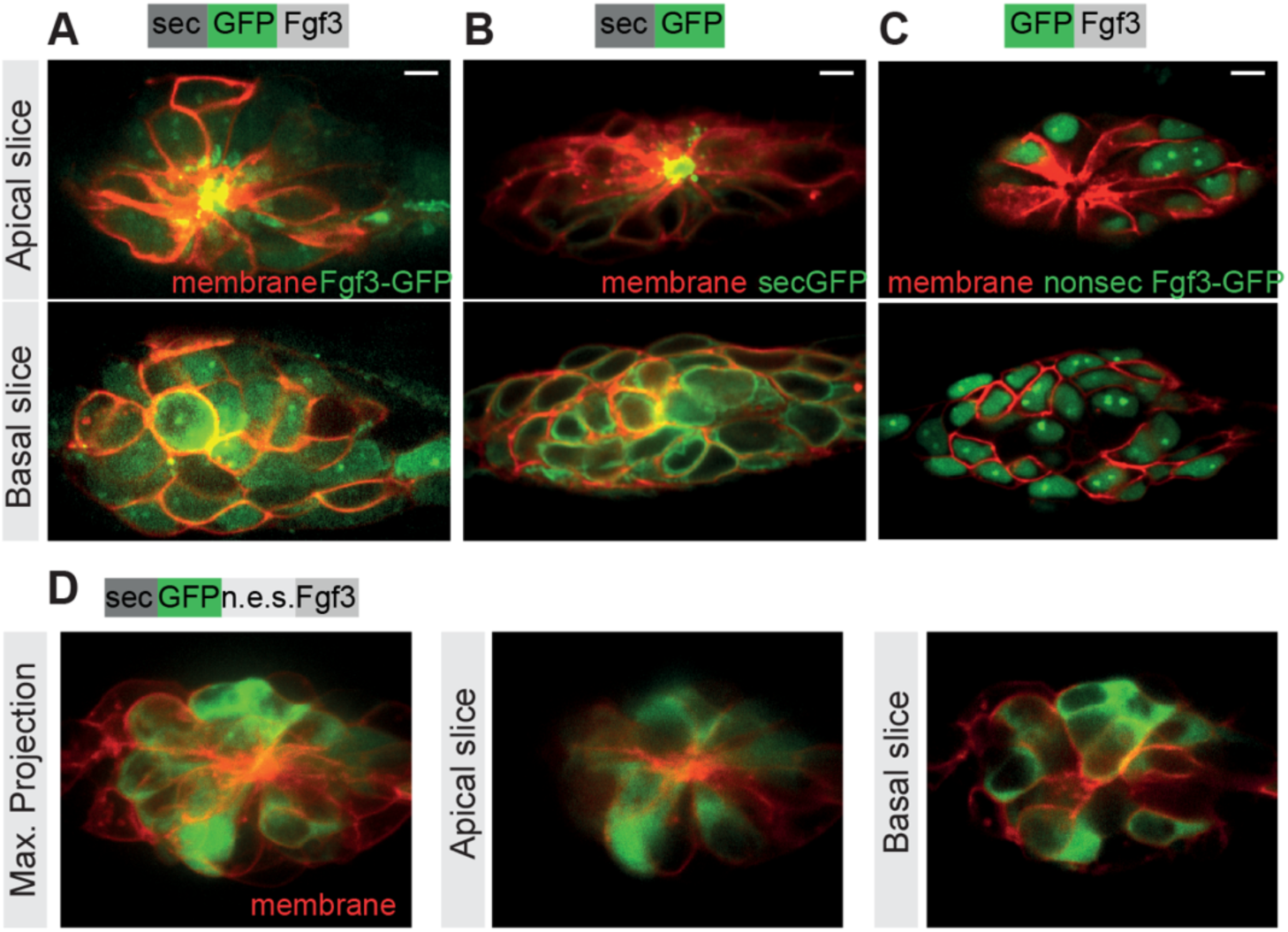
Cellular localization of Fgf3 constructs in lateral line organ progenitors. **A-C.** Apical and basal slices of the imaged organ progenitors as shown in Figure 6A-C, highlighting the corresponding extracellular (apical lumen in rosette center) and cellular localizations within the lateral line organ progenitors. Scale bar: 5 µm. **D.** Apical, basal slices, and the maximal projections of Fgf3-GFP construct with nuclear export signal (n.e.s), showing expected absence of nuclear Fgf3 and unintended block of Fgf3 localization to the extracellular space.

**Figure S9.**
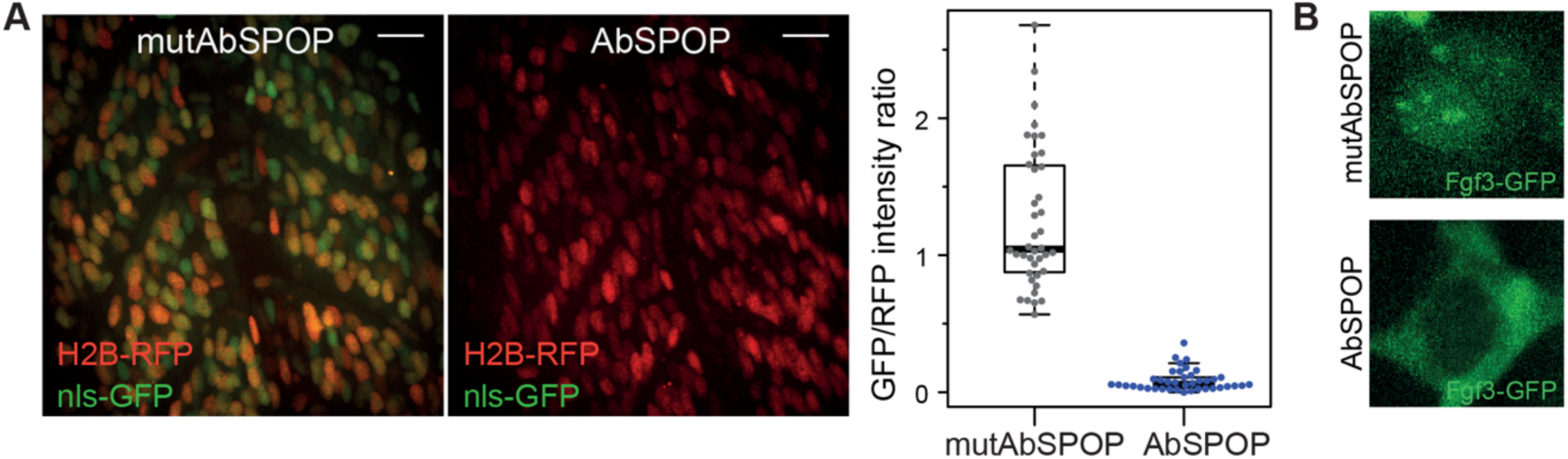
Establishing a nuclear-targeted GFP degradation system for selective depletion of nuclear Fgf3-GFP. **A.** Specificity and efficiency of the anti-GFP degradation tested in zebrafish somites with triple RNA injections of H2B-RFP (red), nls-GFP (green) and either mutAbSPOP or AbSPOP. Left: corresponding example images. Right: intensity quantification. **B.** Fgf3-GFP cellular localization (green channel of Figure 6F-G) upon control (mutAbSPOP) and nuclear degradation (AbSPOP). Scale bars: (A) 20 µm.

## Methods

### Zebrafish handling and stocks

Zebrafish (Danio rerio) strains were maintained as described in Westerfield^54^. Embryos were raised in E3 buffer between 26-30°C. All experiments were conducted on embryos younger than 3 dpf, under the guidelines of the European Commission, Directive 2010/63/EU. All chemical treatments were conducted on embryos dechorionated with pronase. Embryos were anesthetized with 0.01% Tricane for live imaging and prior to dissection or chemical fixation procedures. To prevent pigmentation when required, embryos were treated with 0.002% PTU at 24 hpf. For live imaging, embryos were mounted in 0.8% LMP agarose on glass-bottom dishes (MatTek) and kept at 28°C. The following established mutants and transgenics were used: cxcl12a*^t^*^30516^ from Valentin *et al.* ^35^, cxcr4b:nls-tdTomato from Donà *et al.*^30^, cldnb:lyn-GFP^25^, gal4(ETLGA346) uas:mCherry^55^, atoh:tdTomato^56^, uas:Cxcr7b-GFP^30^, cxcr4b:lexPR/lexop:lyn-RFP, lexop:Fgf3-GFP, lexop: secGFP, fgf3:Fg3-GFP and cxcr4b:cxcr4b-tagRFP from Durdu *et al.*^29^.

### Constructs and Transgenic generation

LexOP and UAS responder lines were generated using MultiSite Gateway cloning (Invitrogen) and the Tol2kit^57^: i) LexOP(p5E)/GFP(pME)/fgf3(p3E), ii) UAS(p5E)/ cxcr7a(pME)/GFP(p3E), iii) LexOP(p5E)/cxcr7b(pME)/GFP(p3E), iv) LexOP(p5E)/ caFgfr1(pME)/RFP(p3E), v)LexOP(p5E)/Mycn(pME)/GFP(p3E). All destination vectors carried SV40poly(A) at the C terminus in the final construct. As a transgenic marker, all clones carried a cmlc2:eGFP cassette at the backbone.

ca-Fgfr1a was cloned using the hs:caFgfr1 construct from Marques *et al.*^58^. FGF family and *cxcr7a* cDNAs were generated from total RNA extracts of zebrafish embryos at 30 hpf. FGF open-reading frames (ORFs) were cloned into CMV plasmids, with GFP inserted downstream of the signal peptide, following the same design used for Fgf3.

twf2b:lyn-mKate2 was generated by inserting lyn-mKate2 in Twf2b BAC clone CH211-14I3, replacing the first exon. cxcr7b:cxcr7b-mKate2 was generated by inserting mKate2 in cxcr7b BAC clone DKEY-96H14 and cxcr7b:lyn-sfGFP was generated by inserting lyn-sfGFP in the *cxcr7b* BAC clone, replacing the first exon. Recombinations were carried out by ET recombineering (Gene Bridges). As a transgenic marker, cry:FP/KanR cassette was inserted into the BAC backbone to drive the FP expression in the eye lens. BACs were purified with Large Construct Kit (Qiagen) and transgenic lines were generated by injecting the constructs at one-cell stage.

### Colorimetric In-Situ Hybridization (ISH)

*pea3*, *fgf3* and *cxcr7 in situ* hybridization (DIG probes, anti-DIG Alkaline Phosphatase coupled antibody, NBT/BCIP substrate at 30°C) were performed as described previously^28,35^. *cxcr7a in situ* hybridization was carried out according to standard protocols with probes generated from *cxcr7a* cDNA. All in-situ probes were hybridized at 65°C. To counter-label the primordium, anti-GFP antibody staining (rabbit-anti GFP primary antibody with Alexa488-anti-rabbit secondary antibody) was performed on cldnb:lyn-GFP transgenic embryos during ISH protocol as in Valentin *et al.*^35^.

### Fluorescence Recovery After Photobleaching

Fluorescence Recovery After Photobleaching (FRAP) experiments were performed using an Ultraview VoX spinning disk microscope equipped with a photokinesis unit and Zeiss ×63 water objective. To monitor fluorescence recovery after photobleach, 5 pre-bleach and 45 post-bleach images were acquired at 30 ms per frame. The bleached region was defined with a strip ROI at the edge of the pool. Mean signal intensity over time was quantified in the bleached region, total pool, and the background to calculate recovery half-times. Measurements were imported into easyFRAP software^59^, and recovery curves were analyzed with full-scale normalization and double-term fitting.

### Single molecule fluorescence *in situ* hybridization

Single molecule fluorescence *in situ* hybridization (smFISH) probes (Custom Stellaris FISH probes, Biosearch Technologies) for *pea3* (*etv4*), *cxcr7b* and *tbx1* were designed using the Stellaris RNA FISH probe designer, comprising at least 40 sequence specific oligonucleotides (20bp each). Probes were conjugated to either Cal Fluor 590 (red) or Quasar 670 (far red) fluorophores. Hybridization, imaging and analysis were performed as described previously^29^.

### Clonal and transient misexpression

Cell transplantation protocols^54^ were used to generate Fgf3-GFP misexpression cell clones. Donor cells isolated from cxcr4b:LexPR, lexOP:lyn-RFP, lexOP:Fgf3-GFP, cxcr4b:nls-tdTomato transgenic embryos were transplanted into cldnb:lyn-GFP transgenic recipient embryos. Expression of Fgf3-GFP was induced with 10 µM RU486 at 24 hpf.

Misexpression clones of caFgfr1 were generated by injection of lexOP:caFgfr1-RFP construct into cxcr4b:LexPR, cldnb:lyn-GFP transgenic embryos at one-cell stage. Expression was induced with 10 µM RU486 treatment at 24 hpf. RNA injections at one-cell stage were used to express H2B-RFP, nls-GFP, AbSPOP^46^ and mutAbSPOP mRNAs, generated with in vitro transcription (Ambion mMessage mMachine Kit) using linearized SP6- or T7-containing plasmids. Transient expression of GFP-tagged Fgf2, Fgf3, Fgf4, Fgf8, Fgf10, Fgf13, Fgf19 and Fgf20 in early embryos was achieved by injecting CMV promoter-driven DNA constructs together with in vitro transcribed H2B-RFP RNA at one-cell stage.

Mouse embryonic stem cells (mESCs) with doxycycline inducible gene expression system (HA36CB1Tet3G) were cultured as described in Durdu et.al., 2025 (DOI: 10.1016/j.molcel.2025.07.005). Briefly, cells were grown at 37°C 7% CO2, seeded on plates coated with 0.2% gelatin and maintained in Dulbecco’s modified Eagle medium (DMEM), supplemented with 15% fetal calf serum (FCS), Glutamax, nonessential amino acids, 0.001% 2-Mercaptoethanol and leukemia inhibitory factor (LIF). Cells were recombined with L1-TRE-FGF3-GFP-1L plasmid (where the tetracycline responsive element (TRE) promoter drives the expression of mouse FGF3, GFP tagged between the secretion signal peptide and the globular FGF domain). mESCs were either maintained in stem cell media (15%FCS+LIF) or differentiated through embryoid body formation (10%FCS+5µM retinoic acid). FGF3-GFP expression was induced with 1 ug/ml doxycycline and cells were imaged at 6-hour post induction.

### Gene expression profiling

Three experimental groups were used for gene expression profiling: (i) wild type (cldnb:lyn-GFP, cxcr4b:nls-tdTomato) incubated with 10 µM RU486, 0.1% DMSO; (ii) FGF inhibition (cldnb:lyn-GFP, cxcr4b:nls-tdTomato) treated with 1 µM SU5402, 0.1% DMSO, 10 µM RU486; (iii) FGF induction (cldnb:lyn-GFP, cxcr4b:nls-tdTomato, cxcr4b:LexPR, lexOP:Fgf3-GFP) incubated with 10 µM RU486, 0.1% DMSO. Embryos at 24 hpf stage were treated with inducers and inhibitors for 6 hours. At least 300 transgenic embryos were used per sample. Embryos were dissected with a scalpel by separating trunks and collected in 2 ml tubes. Embryos were dissociated with FACSmax (Genlantis) and passed through 40 µm cell strainer. Around 20,000 cells were sorted by FACS using GFP, RFP and scatter parameters, with gating determined by the lack of lateral line cell populations in cxcl12a mutant embryos.

Total RNA was extracted with the RNeasy kit (Qiagen) and stored at -80°C. RNA quality was validated, and concentration was measured with Qubit. Samples were processed as described in Affymetrix GeneChip System Protocol and hybridized to Zebrafish Gene ST 1.0 arrays.

### Analysis of the microarray data

Affymetrix Power Tools (APT) was used to pre-process and normalize CEL files generated for each sample. Expression values of ‘meta’ probe sets (transcripts) were summarized with the Robust Microarray Analysis (RMA) approach^60^ as implemented in APT (log2 transformed). Quality control of the samples was initially carried out by checking chips with outlier values of ‘average raw intensity signal’ and ‘mean absolute deviation of the residuals’. Detected above background (DABG) method was applied to estimate whether probe sets were higher than the background distribution. As recommended^61^, ‘main/meta’ probe sets were retained if half or more of the associated probe sets were detected (DABG, *P* value<0.05) in ≥50% of replicates of any condition^62,63^. Out of 40,344 ‘main/meta’ probe sets, 24,720 ‘main/ meta’ probe sets were retained for further analysis. They were annotated with the corresponding gene symbols (21,446 unique) derived from the Affymetrix-ZebGene-1_0-st-v1.na34.zv9.transcript data and through manual curation. Unsupervised hierarchical clustering of the samples (Pearson’s correlation coefficient, average-linkage) was used to examine outliers and batch effects. The limma package^64^ (Smyth, 2004) available from Bioconductor^65^ was used to identify differentially expressed genes for the comparisons of the biological groups while accounting for the batch of the replicate experiments. False discovery rate (FDR) was applied for multiple-test correction^66^. Heat maps representing the fold-changes in figures were generated in R.

### Protein structure models

Protein sequences of all FGF family members were compared by multiple sequence alignment using the T-Coffee server^52^ and visualized with ESPript 3.0 server^53^. For protein structure modeling, protein sequences for Fgf2, Fgf3, Fgf4, Fgf8a, Fgf10a, Fgf13a, Fgf19, and Fgf20a were retrieved from the SMART database^67^, selecting the “Acidic and basic fibroblast growth factor family” domains. Structural models were generated using AlphaFold3^68^ (https://alphafoldserver.com/) with default parameters, and model confidence was assessed via per-residue pLDDT scores. Electrostatic surfaces were computed using APBS^69^ server (https://server.poissonboltzmann.org/) after preprocessing with pdb2pqr^70^ under standard settings. All models were superimposed in PyMOL^71^ (v2.5.0) using the align function, and electrostatic surfaces and structural features were visualized using custom PyMOL scripts.

Classical NLS are characterized by clusters of positively charged amino acids that are typically exposed in an extended loop conformation^72^. Based on this, we focused on a conserved site (motif 4^39^) in Fgf3 with such structural potential. This site was compared across the zebrafish FGF family using multiple sequence alignment and subsequently examined in 3D structural models to assess its charge, surface exposure, and conformation.

### Statistical analysis

All the statistical analysis was performed in R^73^. Comparisons between two groups were conducted using the non-parametric, two-sided Wilcoxon rank-sum test.

Sample sizes (n) and *P* values (*P*) are provided for each experiment and indicated in figures (*\*P <* 0.05,*\*\*P <* 0.01,*\*\*\*P <* 0.001).

Boxplots are displayed as standard box-and-whisker plots indicating the median and interquartile range. For all experiments, individual data points were overlaid on the boxplots as scatter plots using the beeswarm package, enabling direct visualization of the sample variance and distribution. Scatter and line plots were generated with the ggplot2 package^74^ in R.

*P* values and sample sizes:

**Figure 1C**: *n*_WT_ = 4, *n*_FGFRinh_ = 5, *n*_FGFind_ = 6, ∼40 cells per fish, *P*_WT-FGFRinh_ = 0.015, *P*_WT-FGFind_ = 0.0095, *P*_FGFRinh-FGFind_ = 0.051. **Figure 4B**: *pea3* expression: *n*_ctrl_ = 5, *n*_FGF_ = 11, *n*_neigh_ = 6, *P*_ctrl-FGF_ = 4.6 × 10^−4^, *P*_ctrl-neigh_ = 4.3 × 10^−3^, *P*_FGF-neigh_ = 0.88; *cxcr7* expression: *n*_ctrl_ = 10, *n*_FGF_ = 13, *n*_neigh_ = 7, *P*_ctrl-FGF_ = 1.7 × 10^−4^, *P*_ctrl-neigh_ = 0.53, *P*_FGF-neigh_ = 1.2 × 10^−3^. **Figure 5D**: Calculated mobile fraction: nls-GFP 0.98, Fgf3-GFP 0.83, Mycn-GFP 0.86; t_1/2_ (s): nls-GFP 0.24, Fgf3-GFP 0.95, Mycn-GFP 0.97. **Figure 6D**: *n*_ctrl_ = 6, *n*_nonsecFgf3_ = 6, *n*_Fgf3_ = 6, *pea3*: *P*_ctrl-nonsecFgf3_ = 4.6 × 10^−4^; *cxcr7b: P*_ctrl-nonsecFgf3_ = 2.2 × 10^−3^, *P*_ctrl-Fgf3_ = 2.2 × 10^−3^, *P*_nonsecFgf3-Fgf3_ = 2.2 × 10^−3^. **Figure 6H**: *pea3*: *n*_mutAbSPOP_ = 8, *n*_mutAbSPOP_ = 8, *P_Pea3_* = 1; *cxcr7b*: *n*_mutAbSPOP_ = 18, *n*_mutAbSPOP_ = 24, *P_Cxcr7b_* = 1.1 × 10^−8^. **Figure S3E:** *n_cxcr7b_*_-/-_ = 15, *n_cxcr7b_*_-/-uas*Cxcr7a*misexp_ = 14, *P*_(*cxcr7b*-/-)-(*cxcr7b*-/-uas*Cxcr7a*misexp)_ = 2.6 × 10^−8^.

## Notes

https://www.ncbi.nlm.nih.gov/geo/query/acc.cgi?acc=GSE207538

### Competing Interest Statement

The authors have declared no competing interest.

### Summary of Updates

Abstract and discussion rephrased, further supplementary data added.

